# netANOVA: novel graph clustering technique with significance assessment via hierarchical ANOVA

**DOI:** 10.1101/2022.06.28.497741

**Authors:** Diane Duroux, Kristel Van Steen

**Affiliations:** BIO3 - GIGA-R Medical Genomics, University of Liege, Liege, Belgium; BIO3 - Department of Human Genetics, KU Leuven, Leuven, Belgium

## Abstract

Many problems in life sciences can be brought back to a comparison of graphs. Even though a multitude of such techniques exist, often, these assume prior knowledge about the partitioning or the number of clusters and fail to provide statistical significance of observed between-network heterogeneity. Addressing these issues, we developed an unsupervised workflow to identify groups of graphs from reliable network-based statistics. In particular, we first compute the similarity between networks via appropriate distance measures between graphs and use them in an unsupervised hierarchical algorithm to identify classes of similar networks. Then, to determine the optimal number of clusters, we recursively test for distances between two groups of networks. The test itself finds its inspiration in distance-wise ANOVA algorithms. Finally, we assess significance via the permutation of between-object distance matrices. Notably, the approach, which we will call netANOVA, is flexible since users can choose multiple options to adapt to specific contexts and network types. We demonstrate the benefits and pitfalls of our approach via extensive simulations and an application to two real-life datasets. NetANOVA achieved high performance in many simulation scenarios while controlling type I error. On non-synthetic data, comparison against state-of-the-art methods showed that netANOVA is often among the top performers. There are many application fields, including precision medicine, for which identifying disease subtypes via individual-level biological networks improves prevention programs, diagnosis, and disease monitoring.

## 1 Introduction

Subjects or objects can often be linked to systems, and studying the differences between their corresponding system representations is of particular interest to precision medicine. Indeed, examples of systems in biology and medicine include the nervous system, the circulatory system, and the respiratory system. The interactome, which refers to the entire complement of interactions between DNA, RNA, proteins and metabolites within a cell, is another example. Genetic alterations and/or environmental stimuli can change these interactions, making the system and its properties subject-specific. Hence, understanding the controlling mechanisms of biological systems linked to an individual may facilitate disease diagnosis, understanding of disease mechanisms and improve treatment management for patients.

Graphs lend themselves perfectly to visualise systems [31]. A graph always consists of nodes and edges as primary building blocks, no matter the context. Only the characteristics of these elements may differ since they can be labelled, attributed, weighted, directed (see Section 2.1). Nodes are directly measured or synthetic, and their connections can be functional or analytically derived via statistical and machine learning models. Graphs may be studied using relevant substructures, such as hubs, communities [47] and graphlets [46]. Multiple graphs that only differ in their connections can be combined into a multiplex to accumulate information and complete the picture [14, 21, 58]. Sometimes, the term “graph” may be reserved to describe an abstract data structure, whereas the term “network” would refer to a concretisation of a graph. Here, the terms graph and network are interchangeably used.

Machine learning has seeded the development of network analytics [10, 37]. Representation learning [22] is an important subfield of machine learning and is concerned with training algorithms to learn useful representations. In particular, mappings are learned that embed nodes or sub-graphs into a low-dimensional vector space. Structural properties in the embedding should best reflect the corresponding properties of the original graph. The new representations in the embedding space serve as input to downstream analyses, such as classification (supervised learning) or clustering (unsupervised learning). Classification has received the most attention. When entire graphs are embedded, the goal would be to train the algorithms to predict class membership for new graphs. Unfortunately, the majority of unsupervised analyses with graphs involve a single graph only [27]. Yet, as argued before, in the absence of information about class memberships, discriminating between multiple graphs is timely and important [41, 68]. Also, there is a need for efficient unsupervised learning strategies that use graphs with labelled – and thus unexchangeable – nodes as input.

In response to the illustrated shortcomings, with our novel netANOVA analysis workflow we aim to exploit information about structural and dynamical properties of networks to identify significantly different groups of similar networks. We do so by developing a novel group comparison testing workflow that sequentially evolves down a hierarchical tree. The netANOVA test statistic relies on additive partitioning rather than centroids; the latter is typical in traditional Analysis of Variance (ANOVA) hypothesis testing [3]. Statistical significance is assessed empirically to avoid reliance on distributional assumptions. Furthermore, our flexible analysis workflow accommodates small datasets (smaller than 20) as well as larger ones (up to a few thousand), and can be used in multiple contexts via customisable hyperparameter settings, handling weighted, sparse, or multi-layered networks.

In summary, our analysis workflow can be used to identify and formally test for differences between objects that can be represented as graphs. Hence, application areas include, but are not restricted to, precision medicine and the challenging task of identifying endotypes for biomarker development.

## 2 MATERIALS AND METHODS

### 2.1 Network and graphs

A network is a data structure consisting of nodes and edges modelling the relations between two nodes. A network *G* can be defined as *G* = (*V, E*), where *V* is the set of nodes, and *E* are the edges between them. In biology, nodes can be genes, mRNAs, proteins, metabolites, and edges can represent molecular regulation, genetic interactions, co-localisation, or co-occurrence.

For binary networks, a graph is completely described by its adjacency matrix *A* ∈ 0,1_*n*×*n*_, where *A*(*i, j*) = 1 if and only if the link (*i, j*) ∈ *E*. If matrix A is symmetric, then the graph is undirected, otherwise directed. For weighted networks, *A*(*i,j*) = *w_ij_*, with *i,j* ∈ *N*. Attributed networks have labels and/or attributes on the nodes and/or edges. Attributes (resp. labels) are commonly expected to be real values (resp. alphabetic values).

### 2.2 Distances and similarities between networks

Distance and similarity are related concepts: when distance increases, similarity decreases. A ”distance metric” is a function that satisfies the non-negativity, identity, symmetry and triangle inequality properties [43]. Often, some properties are not necessary, and a ”distance measure” may be used. The latter also captures how different two objects are but is a function that does not satisfy at least one of the four properties. A similarity function satisfies the non-negativity, boundedness, identity and symmetry properties. Multiple functions can be used to turn distances into similarities, such as 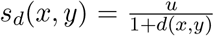, where u is an upper bound, or *d*(*x,y*) = *s_d_*(*x,x*) + *s_d_*(*y,y*) - 2*s_d_*(*x,y*) for kernel-based similarity measures.

The choice of distance measures is a critical step in clustering efforts. An extensive range of graph comparison measures exists. Requiring time-computational efficiency when clustering a large number of graphs dramatically reduces the options. Moreover, most of the remaining distances handle undirected [5, 29, 42] and unweighted [51, 52, 65] networks only. Hence, defining a distance between graphs is a cumbersome task, which requires seeking a context-dependent balance between computational efficiency, performance and interpretability. Following Wills et al. [61], we can group network-based distances into two main classes: Known Node-Correspondence (KNC) and Unknown Node-Correspondence (UNC) methods.

KNC distances depend on the correspondence of nodes. They gather all the methods, such as Euclidean, Jaccard or DeltaCon distances, which require a priori to know the correspondence between the nodes of the compared networks. These methods allow comparing networks where nodes are labelled and hence not exchangeable. In other words, the networks have the same node-set (or at least a common subset), and the pairwise correspondence between nodes is known.

UNC distances gather distances which do not require knowledge of the correspondence between nodes. UNC methods, such as spectral distances, graphlet-based measures, and Portrait Divergence, are suited for global structural comparison. It indicates how much the structures of graphs differ. We will pay special attention to graph kernel measures [8]. A kernel is a measure of similarity between objects and must satisfy two mathematical requirements: it must be symmetric and positive semi-definite. Notably, there are much more UNC approaches than KNC ones.

Our netANOVA workflow accommodates multiple distance measures: edge difference distance [24], a customized KNC version of k-step random walk kernel (see Supplementary) [54], DeltaCon [30], GTOM [67] and the Gaussian kernel on the vectorized networks [16] are proposed as KNC methods. Hamming distance [23], Shortest path kernel [9], k-step random walk kernel, and graph Diffusion Distance [24] are optional UNC methods. More details about these distance and similarity measures, and the reasoning behind these choices are given in Supplementary.

### 2.3 Identification of homogeneous subgroups

Distance-based clustering evolves around finding homogeneous subgroups of objects, where objects with minimal distances between them are assigned to the same cluster. The two most popular distance-based clustering approaches are hierarchical clustering and K-means clustering. The first clusters objects sequentially, via inter-cluster distances. The latter classifies objects into subgroups via inter-cluster variances that need to be minimized. Hierarchical clustering has the additional advantage that a tree (dendrogram) visualizes different granularities in the clustering process, which we will exploit in our workflow.

NetANOVA is built around hierarchical distance-based clustering, with distance measure as in Section 2.2. We use the standard agglomerative clustering which first considers each object as a cluster and then merges clusters successively until one cluster contains all objects.

### 2.4 Deriving the optimal number of clusters

To determine the optimal number of clusters, we recursively test for distances between two groups of networks, progressing from the root node to the end nodes of the clustering dendrogram. Many clustering methods require the user to pre-specify the number of clusters. However, this information is often not known. Incorrect estimation will prevent learning the real clustering structure. Here, the algorithm derives the number of classes. If the two groups created from a node are statistically different, the algorithm to find the optimal number of clusters proceeds in the child nodes. Details about the underlying formal hypothesis test are given next (Section 2.5). There are two stopping conditions: the two subgroups are too small or are not statistically significantly different. The first requires setting a threshold for the minimum allowable size of a subgroup. The result is a decision tree where the end leaves are the final clusters, and splitting is based on a formal group comparison test between network collections. When one of the two groups (a and b) has a size not surpassing the minimum size threshold (for example group a), the statistical test is applied on the other group (group b giving rise to subgroups b1 and b2). If subgroups b1 and b2 are statistically different, group a is regarded to be outlying and hence an independent group.

With the aforementioned sequential procedure, false-positive control is a concern. We include two options to correct for multiple testing. First, we correct the p-values using the depth of the tree, i.e. no correction at the root node, *p_ad_j* = *p* × 2 at level 2 of the dendrogram, *p_adj_* = *p* × 3 at level 3 of the dendrogram, and so forth. Also, we implement the correction developed by Meinshausen [34] and created for variable selection. It controls the FWER at level *α* ∈ (0,1), by performing the hypothesis test described in Section 2.5 at each node j, with the significance threshold: 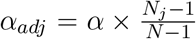 with *N_j_* the number of networks clustered at node j and *N* the total number of networks. It gives increased power to the first nodes (near the root) of the tree. Also, we include the possibility of not correcting for multiple testing in the workflow. Strikingly, computing the total number of tests and applying a Bonferroni correction to each test to keep FWER under control would bypass the hierarchical structure of the analysis.

In Section 3.1 we evaluate these multiple testing corrections for FWER control. We define FWER of the entire workflow as the probability of falsely rejecting the null hypothesis (*H*0: intra variability < inter variability), at least once, when moving down the fixed hierarchical tree, starting from the root node, as explained above.

### 2.5 A novel network-based empirical testing strategy

The netANOVA tests the hypothesis *H*0: intra variability < inter variability, i.e, the variation within a group of graphs is smaller than the variation between groups of graphs.

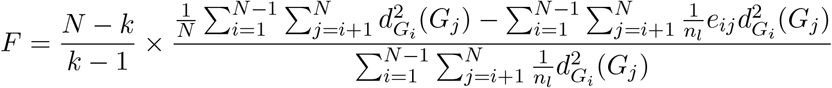

with *N* the total number of individuals, *n_l_* the number of networks in group *l, k* the number of groups, *d_G_i__*(*G_j_*) the distance between graph *i* and graph *j*, and *e_ij_* takes the value 1 if network *i* and network j are in the same group.

A classical ANOVA test [20] uses the mean of a group, but the notion of a mean network is complex. Hence, we take advantage of the following property [3]: the sum of squared distances between points and their centroid is equal to the sum of squared inter point distances divided by the number of points. Another benefit of not using a mean relates to the distance used. For Euclidean distances the mean for each variable across observations within a group constitutes a measure of central location for the group. This is not true for many non-Euclidean distances. The statistic is also interesting in terms of the computational burden. Even though the distance between each pair of networks is required, it is computed only once. No recomputation is required on permutation replicates. In contrast, traditional ANOVA settings require repetitive computation of centroids, network averages and distances to network averages.

Since the actual statistic distribution may not have a closed form and distributional assumptions may not hold on large-sample, significance is derived via permutation replicates. One critical assumption for this test is that the observations need to be exchangeable under a true null hypothesis. Thus, one needs to be careful regarding the interpretation of the significance assessment to ensure that the difference between groups is not due to differences in dispersion (i.e. difference in the distributions). Permutation tests in standard ANOVA settings typically rely on permuting known group labels. In our context, group labels are a priori unknown and inferred via a clustering procedure. Group label reshuffling, conditioning on two clusters in a clustering, will inflate overall type I error [19]. To circumvent this we apply the following procedure to create appropriate null distributions of test statistics. Instead of permuting group labels at each dendrogram node, we permute the distances between the investigated graphs and re-apply hierarchical clustering to identify two groups. If both groups have a size surpassing the group size threshold, we compute the statistics described above. For instance, we repeat the procedure 99 times and compare the permuted statistics to the observed statistics:

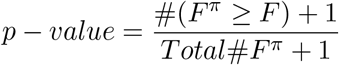

We emphasize that when permuting the values in the original distance matrix, the new matrix cannot be considered a distance matrix because the measure violates the triangular inequality. After applying the permutations, we can indeed obtain *d_G_i__*(*G_l_*) > *d_G_i__*(*G_k_*) + *d_G_k__*(*G_l_*). The linkage criteria in the hierarchical clustering are then limited to methods requiring dissimilarities to be nonnegative and symmetric only, such as complete and average linkage methods [1]. The evaluation of the impact of this linkage criteria is shown in 3.1. In the available code, the user can select ”complete” (default) or ”average” linkage.

In Section 3.1 we also compare different perturbation levels of the distance matrix and set the default amount of perturbation in the distance matrix to 20% and the default number of replicates to 99. These parameters are customisable, as is the significance threshold (default 0.05).

### 2.6 Evaluation and application

All the experiment are conducted on a Scientific Linux release 7.2 (Nitrogen) cluster.

#### 2.6.1 Simulations - Type I error

To evaluate the statistical relevance of the detected groups and the impact of our significance assessment, we study if the proposed workflow controlled the Type I error. We perform simulation analysis based on 1 000 replicates for that purpose. First, we generate an original random graph with m nodes and a density d. For weighted networks, we simulate binary networks and replace the value of edges present by a random number from a normal distribution with a mean 0.5 and standard distribution 0.5*0.5. Edge values are scaled via the min-max scaling algorithm so that values of the adjacency matrix are between 0 and 1. Importantly, we consider the minimum and maximum values across all objects, so these boundaries are the same.

Then, in both the binary and the weighted contexts, we derive n graphs by randomly rewiring the edges while preserving the original graph’s degree distribution [11] of the original graph. Specifically, the algorithm chooses two arbitrary edges in each step (namely (*N_a_,N_b_*) and (*N_c_,N_d_*)) and substitutes them with (*N_a_,N_d_*) and (*N_c_,N_b_*) if they do not yet exist.

We evaluate the impact on the type 1 error of the level of perturbation, the number of graphs, the number of nodes, the graph density, the minimum group size, the distance used to compute dissimilarity between graphs and the minimum number of networks per group. We start from an original baseline network where the original network has a random structure (simulated from Erdos-Renyi model), a density of 0.05, 100 nodes and binary edges. When at least two significant groups are observed in a permutation, that permutation is considered a false positive (FP). This allows us to compute the type I error rate as 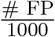.

#### 2.6.2 Simulations - Power

We simulate the situation where each network represents its own individual (e.g., patients), the nodes are labelled and shared across all networks (e.g., genes), and several populations exist (e.g., disease sub-type). The goal is to identify and compare the different populations. To this end, the following experimental setup (see Figure 2.) is implemented. First, we generate an original network and perturb it to derive group networks. Then, we perturb each group network to create individual networks. The goal is to apply the unsupervised netANOVA to assign the individual networks to the correct groups. Finally, we validate the clustering via Jaccard similarity.

**Fig. 1:**
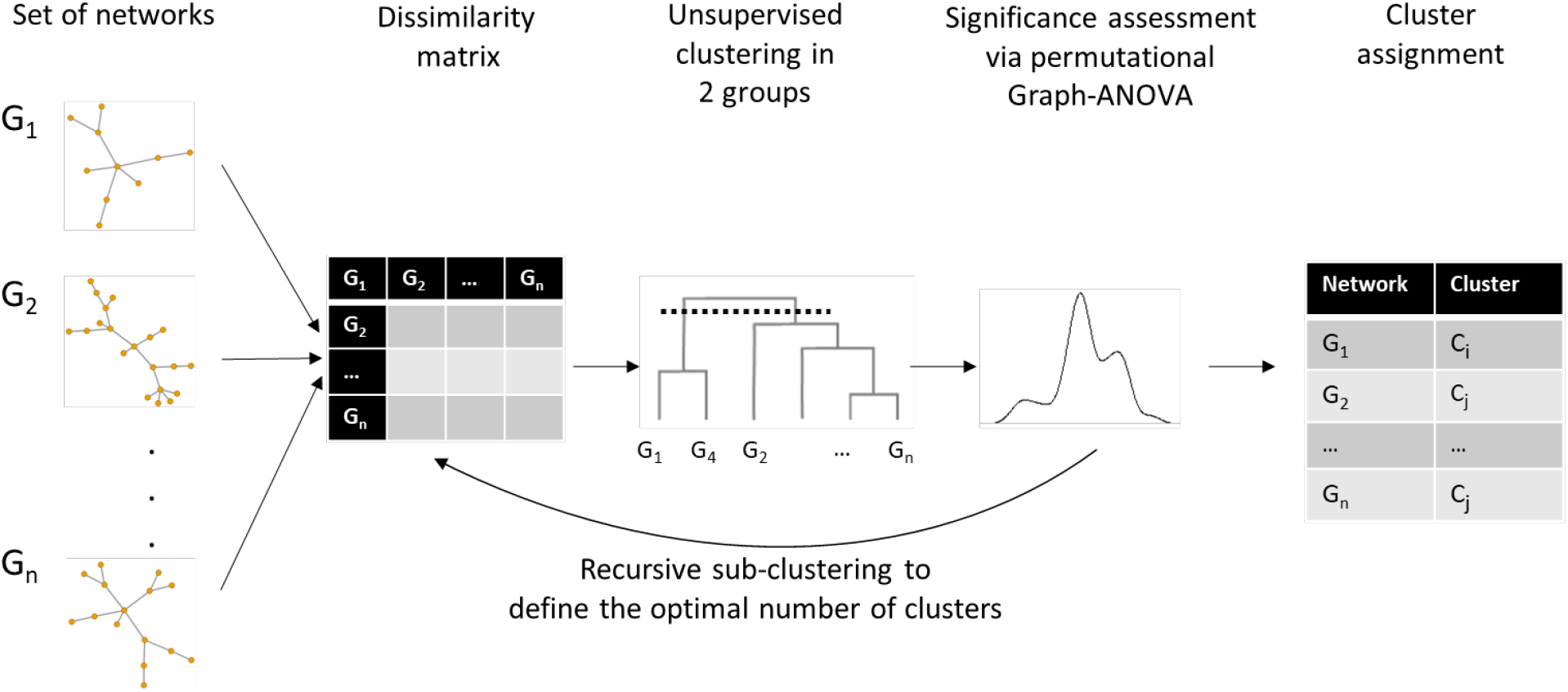
netANOVA workflow

**Fig. 2:**
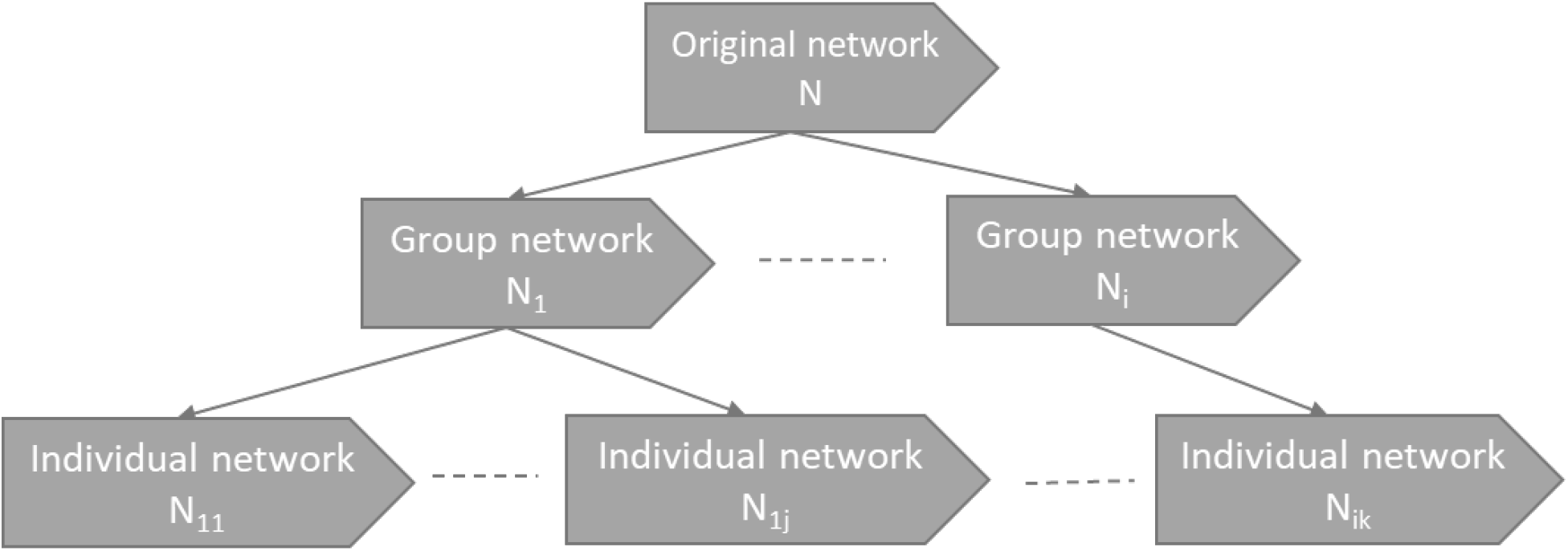
Simulations workflow

We start from the same baseline original networks as in Section type 3.1. First, we randomly switch 40% of the original edges while preserving the degree distribution ten times to create 10 group networks. Then, we switch 40% of the edges for each group network while preserving the degree distribution, 10 times to create 10 individual networks per group.

We create 800 replicates, and we evaluate the impact of multiple parameters. Some of them are associated with the network properties, such as network size or structure. Others are related to the method, such as the correction for multiple testing or the distance between graphs. The influence of the perturbation types and the minimum size of the groups are also studied.

#### 2.6.3 Real-life data application

We apply netANOVA to two real-life bioinformatics graph datasets. For both applications, we use datasets with known clusterings to be able to use these ”true” clusters to compute the performance of netANOVA. Hence, a supervised model could be used to define group membership. Still, the goal here is to evaluate the unsupervised procedure; we do not use any information about the groups in the netANOVA workflow.

##### UNC scenario

The graph dataset MUTAG [13] contains collection of nitroaromatic compounds. The aim is usually to predict the mutagenicity of the compounds on Salmonella typhimurium. The vertices represent atoms, while edges are bonds between the corresponding atoms. The dataset includes 188 samples of chemical compounds. It is publicly available and commonly used to compare classification performances.

##### KNC scenario

We also apply netANOVA to graphs with known node correspondence, i.e. multiplex networks. Previous work [35, 48] has shown the potential of brain networks to distinguish between various brain disorders. We selected the COBRE brain networks [2]. It contains 124 individual-specific networks: 70 controls and 54 schizophrenics. The brain networks are constructed from imaging data (resting state fMRI) to represent functional connectivity between regions of the brain. The graphs are composed of 263 nodes obtained from the Power parcellation [45] and 34,453 edges. The edge weights are the Fisher-transformed correlation between the fMRI time series of the nodes. Nuisance covariates like age, gender, motion (meanFD and meanFDquad) and handedness have been regressed out. For a description of the preprocessing steps to obtain the network edge weights, see [48].

### 3 RESULTS

All adopted simulation and real-life application parameters settings and choices are summarized in Supplementary table 1.

#### 3.1 Type I error

We first investigated the influence of the network properties on the type I error (see Table 1.). Some measures gave rise to a type I error under control in all experimental settings: edge difference, Hamming distance, shortest path kernel, k-step random walk kernel, DeltaCon distance, and gaussian kernel. The graph diffusion distance was more prone to type I error. The network density had a high impact: a higher density gives rise to more conservative results. We also evaluated the algorithm on weighted networks. Although some distances became highly conservative, most of them tended to behave as in the baseline.

**Table 1:**
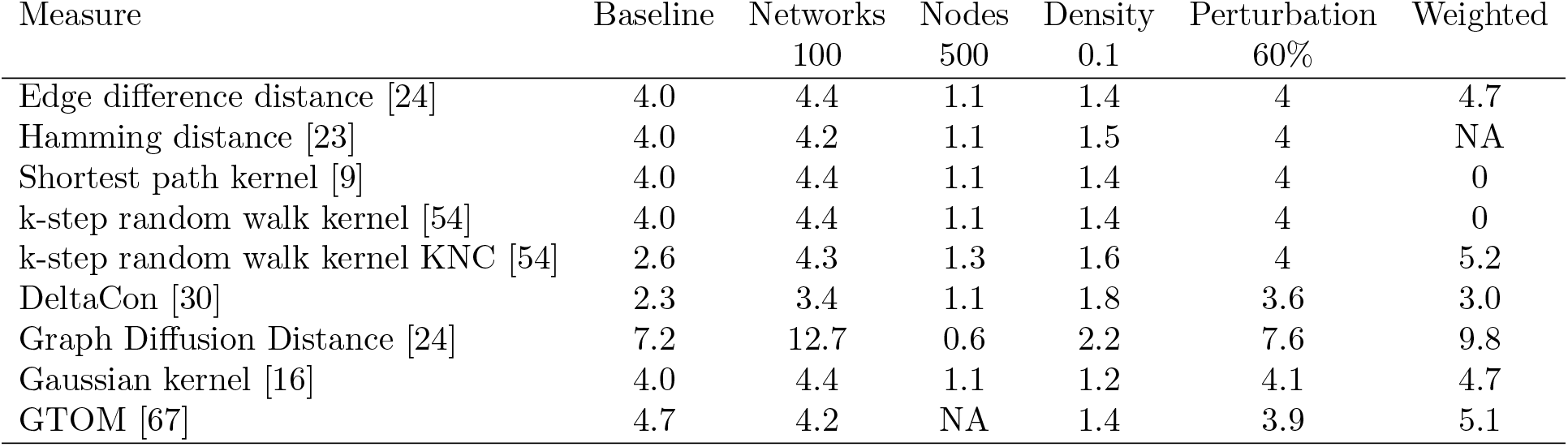
Type I error (%) of the netANOVA workflow depending on network properties, estimated over 1 000 random replicates, as explained in Section 2.6.1. Baseline corresponds to 50 networks, 100 nodes, density 0.05, 40% of the edges are switched, the minimum group size is 10, 20% of the distance matrix is shuffled in the permutations, and the linkage method in the hierarchical clustering is ”complete”.

| Measure | Baseline | Networks | Nodes | Density | Perturbation | Weighted |
| --- | --- | --- | --- | --- | --- | --- |
|  |  | 100 | 500 | 0.1 | 60% |  |
| Edge difference distance [24] | 4.0 | 4.4 | 1.1 | 1.4 | 4 | 4.7 |
| Hamming distance [23] | 4.0 | 4.2 | 1.1 | 1.5 | 4 | NA |
| Shortest path kernel [9] | 4.0 | 4.4 | 1.1 | 1.4 | 4 | 0 |
| k-step random walk kernel [54] | 4.0 | 4.4 | 1.1 | 1.4 | 4 | 0 |
| k-step random walk kernel KNC [54] | 2.6 | 4.3 | 1.3 | 1.6 | 4 | 5.2 |
| DeltaCon [30] | 2.3 | 3.4 | 1.1 | 1.8 | 3.6 | 3.0 |
| Graph Diffusion Distance [24] | 7.2 | 12.7 | 0.6 | 2.2 | 7.6 | 9.8 |
| Gaussian kernel [16] | 4.0 | 4.4 | 1.1 | 1.2 | 4.1 | 4.7 |
| GTOM [67] | 4.7 | 4.2 | NA | 1.4 | 3.9 | 5.1 |

Then, we quantified the impact of the algorithm options (see Table 2.). Overall, the type I error was still under control in almost all settings. The type I error tended to deflate when decreasing the minimum group size. Furthermore, the linkage criterium in the hierarchical clustering significantly impacted the false positive rate. The complete linkage was more appropriate to detect all the groups than the average linkage from the simulations, which is highly stringent. Thus, the complete linkage was set as the default option. Finally, the higher the number of perturbations in the distance matrix in the netANOVA permutation procedure, the more conservative the test.

**Table 2:** Type I error (%) of the netANOVA workflow depending on netANOVA parameters, estimated over 1000 random replicates, as explained in Section 2.6.1 I error. Baseline corresponds to 50 networks, 100 nodes, density 0.05, 40% of the edges are switched, the minimum group size is 10,20% of the distance matrix is shuffle in the permutations, and the linkage method in the hierarchical clustering is “complete”.

| Measure | Min group size<br>5 | Linkage<br>average | Perturbation of<br>distance matrix 10% | Perturbation of<br>distance matrix 50% |
| --- | --- | --- | --- | --- |
| Edge difference distance [24] | 3.5 | 0 | 4.2 | 3.2 |
| Hamming distance [23] | 3.5 | 0.2 | 4.4 | 3.4 |
| Shortest path kernel [9] | 3.5 | 0 | 4.2 | 3.2 |
| k-step random walk kernel [54] | 3.5 | 0 | 4.2 | 3.2 |
| k-step random walk kernel KNC [54] | 2.1 | 0.1 | 2.5 | 1.6 |
| DeltaCon [30] | 2 | 0.2 | 2.2 | 1 |
| Graph Diffusion Distance [24] | 6.2 | 0.9 | 4.7 | 7.5 |
| Gaussian kernel [16] | 3.4 | 0 | 4.4 | 3 |
| GTOM [67] | 4.2 | 0.7 | 4.8 | 4 |

### 3.2 Power

Overall, the baseline scenario (i.e. graphs with random structure, density 0.05, 100 nodes and binary edges) gave good performance with a mean Jaccard index of 0.85 across all distances (see Fig. 3). The correction for multiple testing developed by Meinshausen [34] (see Section 2.4) was less conservative that the correction using the depth of the tree and was, therefore, more optimal with the chosen baseline parameters. Indeed, the former detected nine groups on average across distances versus six groups for the latter. To validate the trends identified in the section 3.1, we applied the average linkage in the hierarchical clustering. Here, it detected only seven groups on average. It confirmed that this linkage is more stringent than the complete one and makes the detection of the correct clusters more complex.

**Fig. 3:**
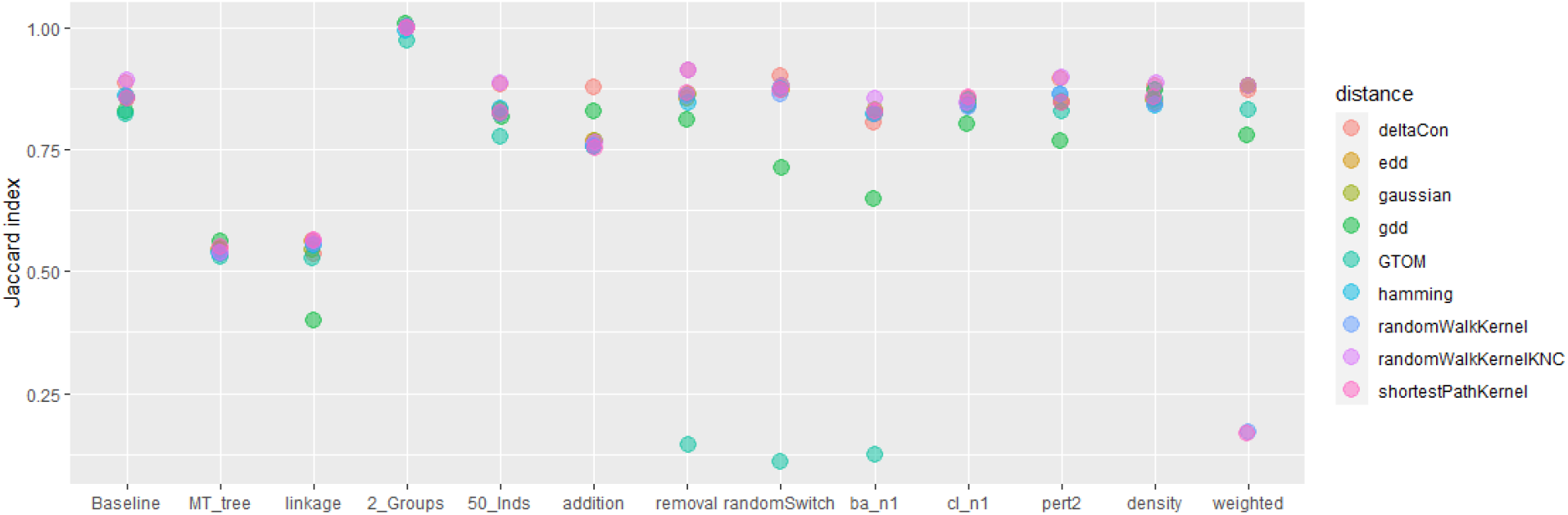
Average Jaccard index across multiple simulation scenario for 1000 replicates. *MC tree* is multiple testing correction using the depth of the tree instead of developed by Meinshausen [34]. *Linkage* stands for average linkage in the hierarchical clustering. *50 Inds* is 50 network per group. Then, *addition* and *removal* correspond to addition and removal of edges instead of rewiring. *Ba_n1* is scale-free networks (from Barabasi-Albert models), *cl_n1* is cluster networks. *Pert2* is associated with an increased within and between group difference of 60% of edge rewiring instead of 40%. *Density* corresponds of a network density of 0.1 instead of 0.05 and *weighted* is for weighted networks (see Section 2.6.2.

We also compared various graph characteristics. We noted that with two groups only instead of 10, the classification was perfect for almost all distances. Also, when we simulated larger groups (50 graphs per cluster), the Jaccard index was comparable to the one obtained with ten networks per group since it ranged between 0.79 and 0.89. Then, we tested multiple perturbation types: random switching, removal of edges and addition of edges. GTOM was less indicated when the perturbation was removal of edges (respectively, random switching) since the associated Jaccard index is 0.14 (resp. 0.1) on average. We also modified the original network structure and tested scale-free graphs using Barabasi-Albert models and cluster networks. with scale-free networks, across all distances except these GTOM and graph diffusion, the average Jaccard index was again relatively high (0.83). The average Jaccard index per distance ranged from 0.79 to 0.86 across all distances with cluster networks.

Moreover, we investigated the impact of perturbations within and between groups of networks by increasing this level up to 60%. The average Jaccard indexes were not highly different from those obtained with the baseline. Then, we increased the density of networks. With a density of 0.1 instead of 0.05, the average Jaccard index ranged from 0.85 to 0.88 and hence, groups were still detectable. We also tested weighted networks (see Section 3.1), and we saw that distances based on random walk kernel didn’t perform as good as the other distances. In most settings, we noted that the graph diffusion distance tended to have difficulties clustering the graphs correctly. On the contrary, DeltaCon and the custom random walk kernel performed overall better than the other measures.

### 3.3 Real-life data application

#### UNC scenario

We compared the results of netANOVA to the recently developed methodologies DGI [59], InfoGraph [55], GraPHmax [7] and its variants (see Table 3). The default options were selected in netANOVA. We derive the distance matrix from the random walk kernel and we set the minimum group size to 40 networks. The algorithm detected the correct number of groups (2) and the associated accuracy is 79.8%. Importantly, in the other methodologies, k-means algorithm (with k=#unique labels) was used on the vector representations of the graphs obtained from the different algorithms. In other words, the number of groups was forced, whereas we didn’t need to specify it with netANOVA. Overall, results showed that netANOVA was able to achieve competitive performance while being able to determine the number of groups and assess statistical significance.

**Table 3:**
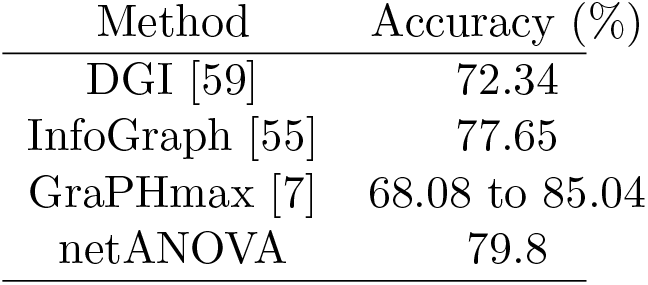
UNC scenario: the clustering accuracy of unsupervised graph-based algorithm

#### KNC scenario

We compared netANOVA performance to six approaches previously used on this dataset: graphclass [48], Elastic net [18], SVM-L1 [69], Signal-subgraph [60], DLDA and LASSO (see Table 4). In netANOVA, the edge difference distance was applied because nodes are labelled and weighted. The minimum group size was set to 10 and the default options were used. Since networks are originally fully connected, we evaluated the impact of graphs sparsification using the method developed by Relión et al. [48] to select edges. When focusing on 6000 or less, a minimum accuracy of 91.9% was obtained. However, when too many edges were considered, we couldn’t distinguish between cases and controls. In the context of brain networks, it was already reported that feature selection is required to detect differences between groups (see Supplementary). When focusing on relevant edges, netANOVA was again among the top performers even compared to supervised methods which usually lead to an inflated accuracy since the phenotype is used in the model. Hence, netANOVA was able to identify groups from a set of networks where nodes are labelled when proper edge selection was performed a priori.

**Table 4:** KNC scenario: accuracy for different methods with variable selection.

|  | Method | Accuracy (%) |
| --- | --- | --- |
| **Supervised**<br>Cross-validated accuracy (average<br>and standard errors over 10 folds) | graphclass [48] | 92.7(2.6) |
|  | Elastic net [18] | 89.5 (1.8) |
|  | SVM-L1 [69] | 87.9 (2.2) |
|  | Signal-subgraph [60] | 86.1 (3.3) |
|  | DLDA | 84.6 (3.3) |
|  | LASSO | 80.1 (5.6) |
| **Unsupervised** |  |  |
| | netANOVA $\rho = 1$ (3,766 edges) | 91.6 |
| | netANOVA $\rho = 0.8$ (4,817 edges) | 100 |
| | netANOVA $\rho = 0.6$ (6,283 edges) | 100 |
| | netANOVA $\rho = 0.4$ (9,606 edges) | no group detected |
| | netANOVA $\rho = 0.2$ (33,796 edges) | no group detected |
| | netANOVA $\rho = 0$ (34,453 edges) | no group detected |

## 4 DISCUSSION

In this article we propose a novel workflow for statistical graph clustering, evaluate its properties (Section 3.1 and 3.2) and validate it via biological networks use cases (Section 3.3 and 3.3). Overall, the type I error is under control. The extensive simulations show that netANOVA can reach high performance, both regarding type I error control and power, and show which option to prioritize depending on the context. The applications on real data reveal that the method achieves competitive results since netANOVA is often among the top approaches. This highlights the method’s potential in real-life situations.

Most of the components in the procedure don’t require a high computing time. The most influential aspects are the number of networks, their density, and the distance chosen to compare the graphs. Distances for graphs with no node correspondence often require a longer processing time. Also, the computations of the first permutation-based significance assessments are the most intense due to the number of graphs compared.

### Novelty of the netANOVA strategy

The workflow differs from generic non-parametric multivariate ANOVA [3] and standard clustering methods in several respects. NetANOVA is a comprehensive graph-specific clustering workflow developed on strong statistics. It starts from a set of networks, derives potential groups, determines the optimal number of groups without the need to set externally the number, and assesses statistical significance while being completely unsupervised. Whereas this can be a great advantage for a user, it makes our workflow difficult to compare to baselines. Indeed, common methods often perform only one part of the analysis and there is a lack and a need for such complete methods. For instance, a common strategy in the absence of graph labels and graph comparative analysis is to generate graph embedding, such as Graph2Vec [39], and AWE [25]. These are fed into downstream models, such as a K-means clustering. However, deriving the optimal number of clusters is often decoupled from the mainstream analysis [7], which is not the case in our proposed workflow. Fraiman et al. [17] outline another strategy. These authors examine network differences between groups with an analysis of variance test explicitly developed for networks. They test whether the mean networks for predetermined groups are the same versus the alternative that at least one group has a deviating average network. Significance is derived by randomly distributing observations across groups in which no subgroup differences are to be expected. In contrast, we do not use the notion of an average network. The reason is that such a notion is not always meaningful. Also, our proposed workflow does not assume knowledge about group formation but identifies relevant partitions on the fly. The permutation procedure is also customized to handle the hierarchical structure of the workflow. In addition, the approach includes components to control for type I error, which improves the confidence in the detected groups. Graph Neural Networks (GNNs) [6] have also become a growing topic for supervised and unsupervised network clustering. GNNs are a class of deep learning methods created to perform inference on data represented by graphs, and they can be used to do graph-level prediction tasks. However, they often suffer from a lack of explainability and a high computational burden, which we address in our analysis workflow.

### Distance and similarity measures

We propose a large range of distances between networks in our workflow but plenty of other measures for network comparison have been developed and were reviewed in depth elsewhere. For instance, in Tantardini et al. [57], the authors evaluate the performance of such methods, carrying out clusterings. They highlight different behaviours and performances between KNC and UNC methods. When networks of the same size and density are considered, most procedures can reasonably discriminate between different structures in the undirected and directed case, often achieving a perfect prediction. However, the results change considerably when considering different sizes and densities of networks. In both the undirected and the directed cases, the graphlet-based measures GCD-11 and DGCD-129 demonstrate superior performance to the other methods in discriminating between different network topologies and outperforming all the other methods investigated. In Wills et al. [61], the authors observe that the adjacency spectral distance exhibits good performance for the comparison graphs of different sizes and the comparison of graphs without known vertex correspondence. Shimada et al. [53] also highlight that graph distance based on the comparison of their Laplacian matrices is interesting because the Laplacian matrix contains essential information about the structural and dynamical properties of networks. In addition, when examining the global structure, Wills et al. [61] find that the adjacency spectral distance and DeltaCon distance both provide good performance. These authors recommend using a spectral distance computed from either the combinatorial graph Laplacian or the adjacency matrix to distinguish graphs via their mesoscale connectivity structures. However, the adjacency spectral distance is not the most appropriate choice in any situation. Wills et al. [61] studied the problem of detecting change points in a dynamic graph. To detect change points, the distance between consecutive time steps is calculated. In that context, the two compared networks share many more edges than in the usual two-sample test. The matrix distances such as the resistance perturbation distance, or DeltaCon, give very high performance and perform better when detecting changes in the network dynamics variables. In the meantime, the spectral distances, yield deficient performance. Therefore, it is crucial to know whether local topological features are of interest in the graph comparison. If local structures are not informative, selecting distances focusing on such structures can harm the accuracy. On the other hand, important information can be lost when local structure is not considered. Hence, the choice of the distance is domain-specific, will depend on the nature of the networks and will vary with the information one is interested in. For instance, the user may initially focus on large-scale network structures (e.g. community structure or the hubs), or small scale features (e.g. local connectivity or triangle graphlets). We recommend to test multiple distance measures and combine the analysis of graph structures to derive a consensus [36].

### Clustering

We selected hierarchical clustering to group networks based on distances, but a multitude of algorithms exist that enable clustering. Here, we discuss the main categories of clustering methods: hierarchical, partitional, grid, density-based, and model-based. Hierarchical clustering forms groups by iteratively dividing or aggregating the objects in a top-down or bottom-up manner. Partitional clustering optimizes some criterion functions such as the Euclidean distance between the object with each of the available clusters. It assigns the object to the closest cluster (k-means [32, 33], fuzzy k-means [66]). Grid clusterings [44] partition the objects into a finite number of cells to create a grid structure and derive groups from these cells. In density-based methods, clusters are separated from other groups other by contiguous regions of low point density (DBSCAN [15, 62], Optics [4]). In mixture density-based methods, objects are assumed to be generated from probability distributions and can be derived from several categories of density functions, or from the same families but with different parameters. Spectral clustering [40] uses information from the eigenvalues of special matrices built from the graph or the data set. Probabilistic clustering (Bayes framework [49]) uses Gibbs posteriors to improve the quantification of uncertainty in the estimated clusters, face computational problems, and large sensitivity to the choice of kernel.

The different methods and their associated aims and assumptions can confuse the user. Many reviews [26, 38, 50, 63, 64] compare approaches. Overall, they underline that no clustering algorithm universally outperforms the others in all contexts. Whereas optimality is often defined in terms of excellent cluster separation and within-cluster homogeneity for an increasing number of applications in biomedicine, the best clustering algorithms will be able to deal with a lot of objects and high-dimensional features. In the meantime, it will be scalable, in terms of storage requirements and running. It should identify irregular shapes of classes and handle outliers and noise. The most relevant methods will also decrease their reliance on parameters that are set by the users. Optimally, the clustering would be able to deal with new data without recomputing the classes from scratch. It will not be dependent on the order of the input patterns. It will handle multiple data types, such as quantitative and qualitative inputs. Also, it should give a reasonable estimation of the final number of clusters without prior knowledge. Finally, it will provide relevant data visualization.

In netANOVA, we chose hierarchical clustering for the following reasons. This algorithm can detect arbitrary cluster shapes rather than being restricted to common shapes. It is insensitive to the order of input patterns and doesn’t rely on a large number of parameters. In addition, whereas we only input continuous values (distances) in our examples, it accepts different data types which makes it easily extendable to different contexts. Strikingly, it provides results as an informative tree structure that is readily interpretable. This visualization helps in understanding and identifying the number of clusters.

### Significance assessment

Several choices were made in the significance assessment procedure. The permutation-based significance assessment can not be performed as in classical non-parametric distance-wise ANOVA [3] because the clusters are derived via hierarchical clustering. Even if there are no actual groups, the clustering will create it by grouping the most similar networks, decreasing the within-group variance and increasing the across-group variance. Thus, a permutation of the graph labels to compute a p-value will bias toward false positives. Since the significance assessment is conditional on the two groups, because of the hierarchical clustering, the same data must not be used to perform clustering and assess significant differences between groups. Multiple suggestions have been suggested to tackle this issue. In Gao et al. [19], the authors propose a selective inference approach to test for a difference in means between two clusters. Kimes et al. [28] developed a Monte Carlo based approach for statistical testing significance in hierarchical clustering. Suzuki and Shimodaira [56] developed the R package pvclust where the hypothesis tests are based on bootstrapping procedures. Our approach also relies on randomisation of the observed data, using permutations of the distances between the investigated graphs instead of the graph label. We re-apply the hierarchical clustering on these permuted sets to identify two groups and compare the obtained labels to the observed ones. Since the permutation of the distance has the additional impact that it no longer satisfies the triangular inequality, the linkage method in the hierarchical clustering is restricted.

### Userfriendliness of netANOVA

There are multiple options in the workflow, such as the distance, the multiple testing correction method, the hierarchical linkage criteria, the minimum size of a group to be tested, the significance threshold, the number of permutations and the percentage of distances permuted in the distance matrix. It can therefore adapt to multiple scenarios and network types. The customisable properties of netANOVA make it relevant to a larger range of users. For example, even though netANOVA has been developed for network analyses, it is generic in that it can accommodate any type of object. The only prerequisite is that a meaningful pairwise distances matrix can be computed.

### Future enhancements

Our netANOVA workflow in the context of high-density networks can be improved. For now, edge selections may be required to select the most informative subnetworks and must be performed a priori. In our KNC application, the edge selection in COBRE networks is supervised and applied on the same dataset as the clustering. Even if the clustering is then performed unsupervised, this could lead to overoptimistic performance estimates. This KNC application shows the importance of focusing on relevant interactions to improve interpretability and accuracy. Thresholding is typically adopted to cancel a percentage of the weakest connection, to turn fully connected and weighted brain networks into a useful sparse network. De Vico Fallani et al. [12] indicate that the way to fix this threshold is still an open issue, and they introduce a criterion, the efficiency cost optimisation (ECO), to select a threshold based on the optimisation of the trade-off between the efficiency of a network and its wiring cost. It is essential to filter out noise, i.e. non-informative elements in the graphs. ”Informative” parts can be extracted in non-supervised ways [14] or with more specific methods, for instance looking for areas in the networks that exhibit a lot of variation between individuals, assuming that the more variation we have in ”the input”, the more we will be able to explain with it. On the other hand, for weighted networks, even when we have a selection of nodes under consideration, the network will still be dense. Hence, some approaches based on multiple thresholds, such as filtration curves can be considered to capture a balance between hard thresholding and fully connected networks. The different thresholds reveal different structures in the graphs, and how these structures change from one threshold to another may be quite different from one network to another.

## 5 CONCLUSION

This study presents a novel workflow for statistical comparison of networks. Extensive simulations showed that netANOVA could achieve high performance in many scenarios while controlling type I error. We also used non-synthetic data to compare its performance against standard and state of the art methods, and we observed that netANOVA is often among the best performers.

Our netANOVA workflow is different from conventional approaches in many aspects. First, it involves reliable statistics that consider the specificities of graphs to enhance our belief in the discovered groups of networks. Second, it performs a comprehensive graph comparison that doesn’t assume knowledge about the partitioning and includes deriving the optimal number of classes. Third, it is user-friendly and flexible because users can customise multiple options to adapt to specific contexts and network types. For example, there is a trend to describe patients via individual-level biological networks in network medicine. NetANOVA facilitates the identification of relevant disease subtypes and endotypes, or strata for adapted disease management, in this scenario. We believe that netANOVA can have a high impact on precision medicine.

## 6 Data Availability

The code necessary to reproduce this article’s results and analyses is available on GitHub at https://github.com/DianeDuroux/netANOVA.

## 8 ACKNOWLEDGEMENTS

We are grateful to Karsten Borgwardt and Leslie O’Bray for their advice and interesting discussions.

