## Supplementary for "netANOVA: novel graph clustering technique with significance assessment via hierarchical ANOVA"

### netANOVA

#### SUPPLEMENTARY DATA

##### Details on distances included in the workflow

Our workflow proposes multiple distances and similarity measures with different properties to allow the user to choose the one adapted to its specific use case. We included the simple edge difference distance [5] that inputs two adjacency matrices and takes the Frobenius norm of their differences as a baseline because this method is computationally fast and easy to interpret. In the same vein, we propose the Gaussian kernel [3] that is applied to the vectorised edge weights. It is defined as $k_{gaussian}(x_i, x_j) = exp(\frac{-||x_i - x_j||^2}{\sigma^2})$ where $||x_i - x_j||$ is the Euclidean distance and $\sigma^2$ is the bandwidth of the kernel. Both methods can be applied to weighted and KNC graphs. The hamming distance [4] that counts the discrepancy between two networks for each edge can also be chosen and used on binary networks. Furthermore, the user can select the shortest path kernel [1], which defines the similarity between two graphs in terms of the similarities of their shortest paths. It is the only measure proposed that is based on substructures. It is computationally intensive when the number of nodes increases. Then, we include the k-step random walk kernel [11]. It measures graph similarity by counting matching walks in two graphs. Longer walks of length k are down-weighted by a factor of $\lambda^k (\lambda < 1)$ to ensure convergence of the corresponding geometric series. It is the only method that is based on walks. Both shortest-path kernel and k-step random walk kernel can be applied on directed networks. For a specific situation where a user considers networks having the same set of fully connected nodes, we developed a customised version of the k-step random walk kernel. In that context, only node and edge attributes change from one network to another, and in fact, the graphs are represented in a feature space that is described as a graph, but it is always the same network, so one is comparing different labelling of the same graph. Hence, the random walk kernel is customised with the constraint that the gene ids have to be identical. $k_{RW}$ is defined as follow: $k_{RW}(A_i, A_j) = \sum_{k=0}^{n} \mu(k) q^T W^k p$, with W the Hadamard product (instead of Kronecker product) of $A_i$ and $A_j$ adjacency matrices of individuals i and j, p and q the initial and stopping probability distributions, and $\mu$ a chosen non-negative coefficient. The probabilities p and q can be vectors of 1s to compare individuals through their individual edge weights only; they can also take the node values to compare individuals through their individual edge weights and individual node weights. Another distance that a user can apply is DeltaCon [7]. The first step of the method is

**Table 1:** Parameter choices in simulations and real life applications. In bold, default parameters of the R function.

| Parameter | Type I error | Power | Real life KNC | Real life UNC |
|---|---|---|---|---|
| # Networks | 50-100 | 100-500 | 188 | 124 |
| Density | 0.05-0.1 | 0.05-0.1 | 0.08 to 0.22 | 0.11 to 1 |
| Network type | random | random, scale-free, cluster | NA | NA |
| #Nodes | 100-500 | 100 | 10 to 28 | 263 |
| Edge type | Weigted, unweighted | Weighted, unweighted | Unweighted | Weighted |
| # Groups | 0 | 2-10 | 2 | 2 |
| Distance | All | All | Random Walk kernel | Edge difference |
| Multiple testing | NA | **tree**-Meinshausen | **tree** | **tree** |
| HC criteria | **Complete**-Average | **Complete**-Average | **Complete** | **Complete** |
| Minimum group size | 5-10 | 5 | 40 | 10 |
| Significance threshold | **0.05** | **0.05** | **0.05** | **0.05** |
| # permutations | **99** | **99** | **99** | **99** |
| % Distances permuted | 10-**20**-50 | **20** | **20** | **20** |

to compute the pairwise node affinities via Fast Belief Propagation (FABP) in the two graphs and measure the differences in the corresponding node affinity scores as their similarity score using the root euclidean distance. It computes graph similarity with known node correspondence, and this method is especially useful in detecting changes in the connectivity of graphs. Also, the Graph Diffusion Distances [5] is accommodated. It quantifies the difference between two graphs of the same size and is based on measuring the average similarity of heat diffusion on each graph. It takes an idea from the heat diffusion process on graphs via graph Laplacian exponential kernel matrices and can be applied to the weighted network. A benefit of Graph Diffusion Distances is that it is a metric in the strict mathematical sense. Finally, the option GTOM [14] can be selected. The measure is constructed by counting the number of m-step neighbours that a pair of nodes share and then normalised suitably. It computes a dissimilarity measure based on the notion of topological overlap. Notably, it is independent of the number of paths and the number of geodesic paths connecting a node and its m-step neighbour. Thus, the range of proposed distances allows the user to select the most relevant one depending on the specificity of its input graphs and the graph properties that he wants to capture and that can be context-specific.

#### Comparison of UNC and KNC inputs and outcomes

The two real-life data application settings are very different. In the MUTAG dataset, networks are smaller with about 28 nodes, and categorical edges are recorded with no noise. Graph clustering looks for discriminative patterns, such as communities or paths. This is computationally feasible only on small ones graphs. We observe that the application of netANOVA on this set of networks with exchangeable nodes (see Section 3.3) gives rise to competitive properties. In the context of graphs without exchangeable nodes (see Section 3.3), networks are larger with 263 nodes. The graphs are initially fully connected, edges are weighted and may contain noise since they are statistically derived. We found that there is a need to focus on relevant communities or edges to achieve high clustering performance. Wills et al. [13] come to the same conclusion. They compare sets of connectomes in two types of analysis: weighted connectomes and unweighted connectomes. They vary the density of edges using two thresholds to set functional connectivity. They observe that the variability in the control population is greater than the contrast between the ASD/controls populations, and hence no tested distance separates the two ensembles of connectomes. They highlight that the first reason is that only a subset of edges represent the structural differences between the two graph groups so that the dissimilarities cannot be identified if one uses all the edges. The second reason is that in the studied connectomes, the local changes in connectivity are of the same order of magnitude as the random local variations. Signal-to-noise is, in fact, a recurrent issue in analysing real-life graph data, and particularly in the context of connectivity networks of human brain activity [2]. In the presence of noise, many metrics cannot detect subtle structural differences. We note that others have reported similar findings [6, 9].

##### Brain network analyzes

In brain networks studies, two main approaches are used to compare the graphs. The first one is based on summary measures representing graph topology and ignoring edge weights. It reduces the network to global summary statistics, for instance the average degree, clustering coefficient, or average path length, and use them as new variables. Previous studies [8, 12] have reported significant differences on this type of summary measures for groups of patients with brain diseases compared with controls. Nevertheless, this procedure ignores local structures and can't distinguish local dissimilarities which may decrease the performance of the clustering. The second approach is to consider all edge weights as a long vector, ignoring the network structure. This vector can be inputted into many existing clustering methods. These methods can perform well but it cannot account for network structure either. When feature selection is applied, it can give interpretability at the edge level, but it is often less relevant than identifying differentiating nodes or communities. Instead, we recommend to use feature selection methods derived specifically for graph. For instance, in Arroyo-Relión et al. [10], the authors developed a method that incorporates the network nature of the data via penalties to promote sparsity in the number of nodes, in addition to sparsity penalties that encourage selection of edges. We also recommend to apply KNC methods to study dissimilarities between brain networks such as edge difference distance, DeltaCon or GTOM [5, 7, 14].
